# A transient amphipathic helix in PCSK9’s prodomain facilitates low-density lipoprotein binding

**DOI:** 10.1101/743195

**Authors:** Samantha K. Sarkar, Alexander C.Y. Foo, Angela Matyas, Tanja Kosenko, Natalie K. Goto, Ariela Vergara-Jaque, Thomas A. Lagace

## Abstract

Proprotein convertase subtilisin/kexin type-9 (PCSK9) is a ligand of low-density lipoprotein receptor (LDLR) that promotes LDLR degradation in late endosomes/lysosomes. In human plasma, 30-40% of PCSK9 is bound to LDL particles; however, the physiological significance of this interaction remains unknown. LDL binding *in vitro* requires a disordered N-terminal region in PCSK9’s prodomain. Here we report that peptides corresponding to a predicted amphipathic α-helix in the prodomain N-terminus adopted helical structure in a membrane-mimetic environment; this effect was greatly enhanced by an R46L substitution representing an athero-protective *PCSK9* loss-of-function mutation. A helix-disrupting proline substitution within the putative α-helical motif in full-length PCSK9 lowered LDL binding affinity >5-fold. Modeling studies suggested the transient α-helix aligns multiple polar residues to interact with positive-charged residues in the C-terminal domain. Gain-of-function *PCSK9* mutations associated with familial hypercholesterolemia (FH) and clustered at the predicted interdomain interface (R469W, R496W, F515L) inhibited LDL binding, which was abolished for the R496W variant. These studies inform on allosteric conformational changes in PCSK9 required for high-affinity binding to LDL particles. Moreover, we report the initial identification of FH-associated mutations that diminish the ability of PCSK9 to bind LDL, supporting that LDL association in the circulation inhibits PCSK9 activity.

---

Circulating liver-derived proprotein convertase subtilisin/kexin type-9 (PCSK9) is a leading drug target in cardiovascular medicine due to its ability to bind and mediate degradation of hepatic LDL receptor (LDLR), the primary conduit for the clearance of plasma LDL-cholesterol (LDL-C) (1-3). Gain-of-function (GOF) mutations in *PCSK9* result in familial hypercholesterolemia (FH), whereas loss-of-function (LOF) mutations are associated with life-long reductions in plasma LDL-C and significant protection from cardiovascular heart disease (4-6). Therapeutic monoclonal antibodies that target PCSK9 and prevent its binding to LDLR lower LDL-C by up to 70% in hypercholesterolemic patients, clearly establishing circulating PCSK9 as a central regulator of hepatic LDLR expression and plasma LDL-C levels (7,8).

PCSK9 is a member of the mammalian proprotein convertase family of serine proteases related to bacterial subtilisin and yeast kexin (9). Human PCSK9 is a 692-residue secreted protein consisting of a 30-residue signal sequence followed by a prodomain, a subtilisin-like catalytic domain and a C-terminal cysteine-histidine rich (CHR) domain (Figure 1A). Auto-catalytic processing in the endoplasmic reticulum releases the prodomain, which remains non-covalently bound and blocks substrate access to the active site, rendering mature PCSK9 catalytically inert (10,11). Like LDL, plasma PCSK9 clearance is predominantly via LDLR-mediated endocytosis in liver (12,13). The apolipoprotein (apo) B100 moiety of LDL binds to the ligand-binding domain of LDLR and undergoes acid-dependent release in early endosomes, allowing LDLR to recycle to the cell surface (14,15). In contrast, PCSK9 binds to the first epidermal growth factor (EGF)-like repeat (EGF-A) in the EGF precursor homology domain of LDLR and with higher affinity at acidic pH (16); thus, PCSK9 fails to release in early endosomes and directs LDLR for degradation in late endosomes/lysosomes (3,10,17).

**Figure 1.**
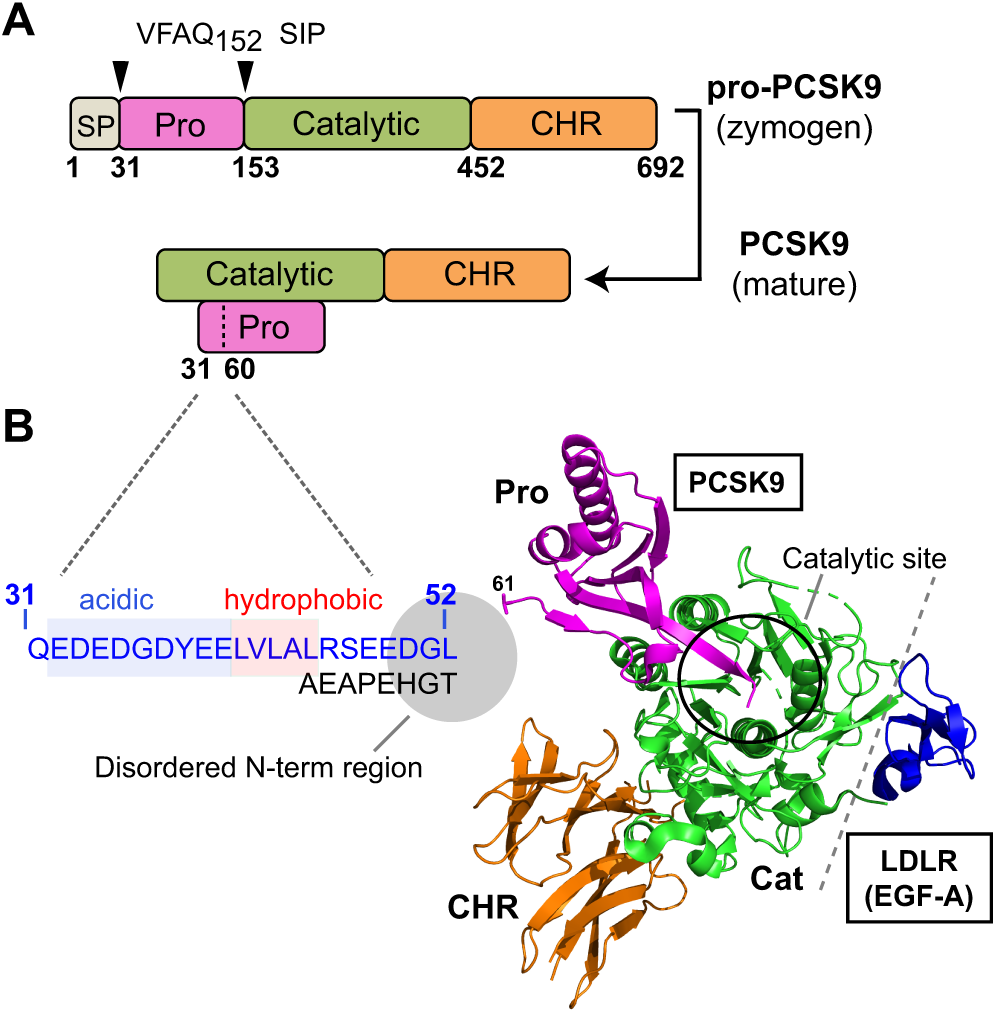
PCSK9 structure with emphasis on disordered N-terminal region of the prodomain. (**A**) Following removal of a signal peptide (SP: aa 1-30, grey) human pro-PCSK9 undergoes autocatalytic cleavage after Gln-152 resulting in mature PCSK9 consisting of a prodomain (aa 31-152, magenta), catalytic domain (aa 153-451, green) and C-terminal histidine-cysteine rich (CHR) domain (aa 452-692, orange). (**B**) Crystal structure of PCSK9 in complex with the EGF-A domain of LDLR (Kwon *et al*, 2008; PBD 3BPS). The C-terminal end of the cleaved prodomain blocks the catalytic site (black circle) which is >20Å from the binding interface with EGF-A (grey dashed line). An intrinsically disordered region (IDR) in the N-terminus of the prodomain (aa 31-60) (shaded circle) is structurally disordered and unobserved in all PDB deposited crystal structures of PCSK9. Highlighted in *blue* is amino acid sequence of an N-terminal region (aa 31-52) required for binding to LDL particles (18). Sequences of interest within this region are a highly-acidic tract (shaded blue) and adjacent hydrophobic region (shaded red).

Approximately 30-40% of PCSK9 in normolipidemic human plasma is bound to LDL particles (13,18). Co-immunopreciptation experiments have confirmed a protein-protein interaction between PCSK9 and apoB100, the main protein component of human LDL (18,19) and PCSK9 has also been shown to bind apoB within the secretory pathway in hepatocytes (20). PCSK9-LDL binding *in vitro* is saturable and specific with a *K*_D_ of ∼125-350 nM (18,21), which is within a range of affinities reported for the PCSK9-LDLR interaction (11,22). Several studies have shown that LDL lowers PCSK9’s ability to bind and mediate degradation of LDLRs in cultured cells (18,22,23). Conversely, there is evidence that LDL association promotes PCSK9-mediated LDLR degradation by inducing a more potent oligomeric form (13,24) or by shielding PCSK9 from inactivating furin-mediated proteolysis (25). In sum, both the molecular mechanism of PCSK9-LDL binding and the physiological significance remain undefined.

We have previously mapped critical LDL binding determinants *in vitro* to an intrinsically disordered region (IDR) in the N-terminus of the PCSK9 prodomain (18). This region has also been identified as a negative allosteric effector of LDLR binding affinity (26,27). A recent study demonstrated the existence of structural flexibility in the prodomain IDR whereby a monoclonal antibody preferentially bound to a transient α-helix (28). Herein, we provide direct evidence demonstrating a functional role of such transient helical conformation in PCSK9-LDL association. Furthermore, computational modeling indicated an intramolecular interaction between the CHR domain and helical conformation of the prodomain IDR. This prompted an assessment of natural *PCSK9* mutations at or near this predicted interdomain interface. Our analysis revealed several FH-associated mutations in the CHR domain that greatly diminished (R469W, F515L) or abolished (R496W) the ability of PCSK9 to bind LDL *in vitro*. These data provide molecular insight on disease-causing mutations in the pathogenesis of FH and support that LDL association exerts an overall inhibitory effect on circulating PCSK9 activity.

## RESULTS

### Identification of a putative amphipathic helix in N-terminus of PCSK9

Figure 1B shows the crystal structure of PCSK9 in complex with the EGF-A domain of LDLR (26) with emphasis on an IDR in the N-terminus of the prodomain (aa 31-60 following the signal peptide cleavage site) that is unresolved in all available x-ray crystal structures of PCSK9 (*e.g.* Refs. 11,29). We have previously mapped important LDL binding determinants to the N-terminal 21 amino acids in the IDR (18). Two sequences of interest are a highly acidic tract (aa 32-40; EDEDGDYEE) and an adjacent hydrophobic segment (aa 41-45; LVLAL) (Figure 1B) both of which are evolutionarily well-conserved in mammals as well as other vertebrates (30) (Supplemental Figure S1). To test potential roles in LDL binding, plasmids encoding human PCSK9 with a deletion of residues 33-40 (leaving Gln-31 and Glu-32 intact for signal peptide cleavage) or replacement of the hydrophobic stretch and a neighboring Arg-46 residue with a glycine-serine linker were constructed and transfected into HEK293 cells. Secreted wild-type (WT) and mutant PCSK9 proteins present in the conditioned medium were then incubated with isolated human LDL particles followed by Optiprep density gradient ultracentrifugation to separate LDL-bound and unbound PCSK9. Deletion of the acidic tract (Δ33-40) nearly abolished PCSK9-LDL binding while replacement of the hydrophobic residues with the Gly/Ser 41-46 linker reduced binding by >60%, suggestive of a functional overlap between these sequences (Figure 2A and B). As recently reported (28), secondary structure predictions using PSIPRED (31) showed an α-helical structure in this region with highest confidence for residues 38-45 (YEELVLAL) overlapping the acidic and hydrophobic segments (Figures 2C). Helical wheel modeling revealed characteristics of an amphipathic α-helix (AH) for these residues, with a relatively polar face and an opposing face consisting of aromatic (Tyr-38) and nonpolar amino acids (Leu-41, Val-42 and Leu-45), giving the helix a directional hydrophobic moment (Figure 2D) (32).

**Figure 2.**
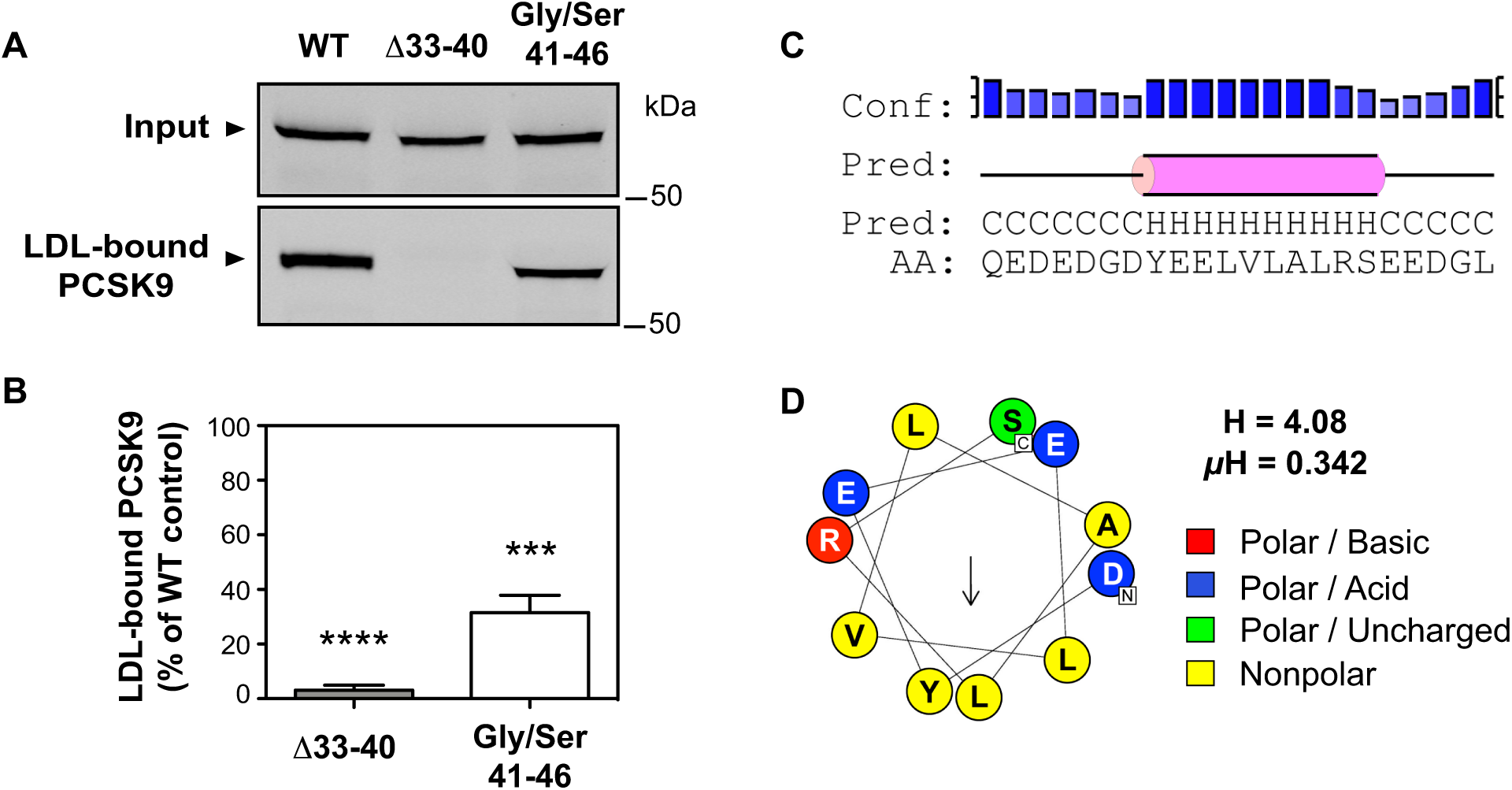
The PCSK9 N-terminal region is predicted to harbor an amphipathic α-helix. (A) Analysis of in vitro PCSK9-LDL binding reactions. Conditioned medium containing wild-type PCSK9 (WT) or variants lacking N-terminal acidic (Δ33-40) or hydrophobic (Gly/Ser 41-46) segments were incubated with LDL prior to density gradient-ultracentrifugation to isolate an LDL fraction and visualization of bound PCSK9 by western blot. (B) Densitometric analyses of western blot in A. Error bars represent SEM (n=5). Significant change in LDL binding compared to WT-PCSK9 control (set to 100%) was determined by One sample t-test: ***, p < 0.001; ****, p < 0.0001. (C) Secondary structure prediction of PCSK9 aa 31-52 by PSIPRED. The prediction (Pred) shows helices (H) as pink rods and random coil (C) as black line. The confidence level (Conf) is shown on the top. Higher bars with darker shades of blue represent higher prediction confidence. (D) helical wheel representation of amphipathic α-helix (residues 37-47) generated from HeliQuest. The arrow indicates the magnitude and direction of the hydrophobic moment (32). The hydrophobicity (H) and hydrophobic moment (μH) from HeliQuest are also shown.

### Amphipathic α-helix formation in a hydrophobic membrane-mimetic environment

Since the putative N-terminal helix is predicted to have amphipathic characteristics, a random coil-to-helix conformation change may be triggered in an amphipathic environment of a membrane surface, such as that present on lipoproteins. Therefore, we performed assessments of the secondary structure that this motif forms in aqueous versus membrane-mimetic conditions. Synthesized peptides corresponding to residues 37-47 (DYEELVLALRS) were dissolved in phosphate buffer or in buffer containing n-dodecyl-phosphocholine (DPC) micelles, a widely used membrane-mimetic that can induce α-helical conformation in peptides with inherent amphipathic properties (33). Circular dichroism (CD) spectra revealed that the motif remained in a random coil conformation in aqueous buffer, but in the presence of DPC micelles adopted a distinctly helical conformation (Figure 3A). Proline is a potential helix-breaker due to a combination of steric constraints and the inability of the backbone imino group to participate in hydrogen bond formation. Accordingly, an A44P substitution in a prodomain-derivative peptide prevented a shift to helical conformation in the presence of DPC micelles (Figure 3B).

**Figure 3.**
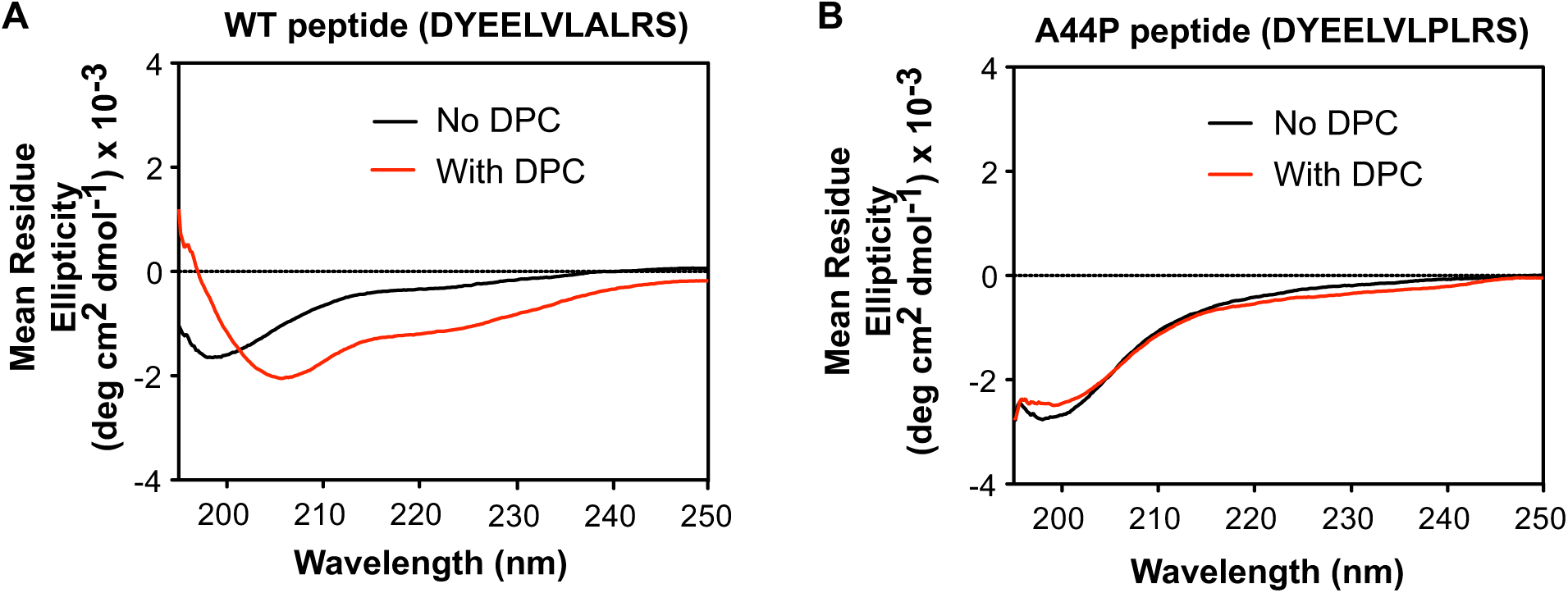
Amphipathic helix formation in the presence of a membrane-mimetic. (A) Circular dichroism (CD) spectra of synthetic peptide representing WT PCSK9 residues 37-47 (DYEELVLALRS) in absence (*black line*) or presence (*red line*) of *n-*dodecylphosphocholine (DPC). Spectra shown are representative of three independent experiments. (B) CD spectra obtained as in *A* using an A44P mutant peptide (DYEELVLPLRS).

### Proline substitutions in the putative N-terminal helix greatly diminish PCSK9-LDL binding

To test the influence of peptide backbone conformation on LDL binding we introduced proline residues at two separate positions in the putative helix (L41P and A44P) in full-length recombinant PCSK9. In each case, proline substitution inhibited the ability of secreted PCSK9 to bind to LDL by >90% (Figures 4A and B), supporting an important structural role of helical conformation in the prodomain N-terminus. Both PCSK9 proline variants were capable of binding to LDLR, as determined by cell uptake assays conducted in HEK293 cells overexpressing human LDLR (Figures 4C and D). In subsequent experiments, we focused on functional effects of the A44P mutation in PCSK9 since the methyl side-chain of alanine is non-reactive and less likely to be directly involved in protein function.

**Figure 4.**
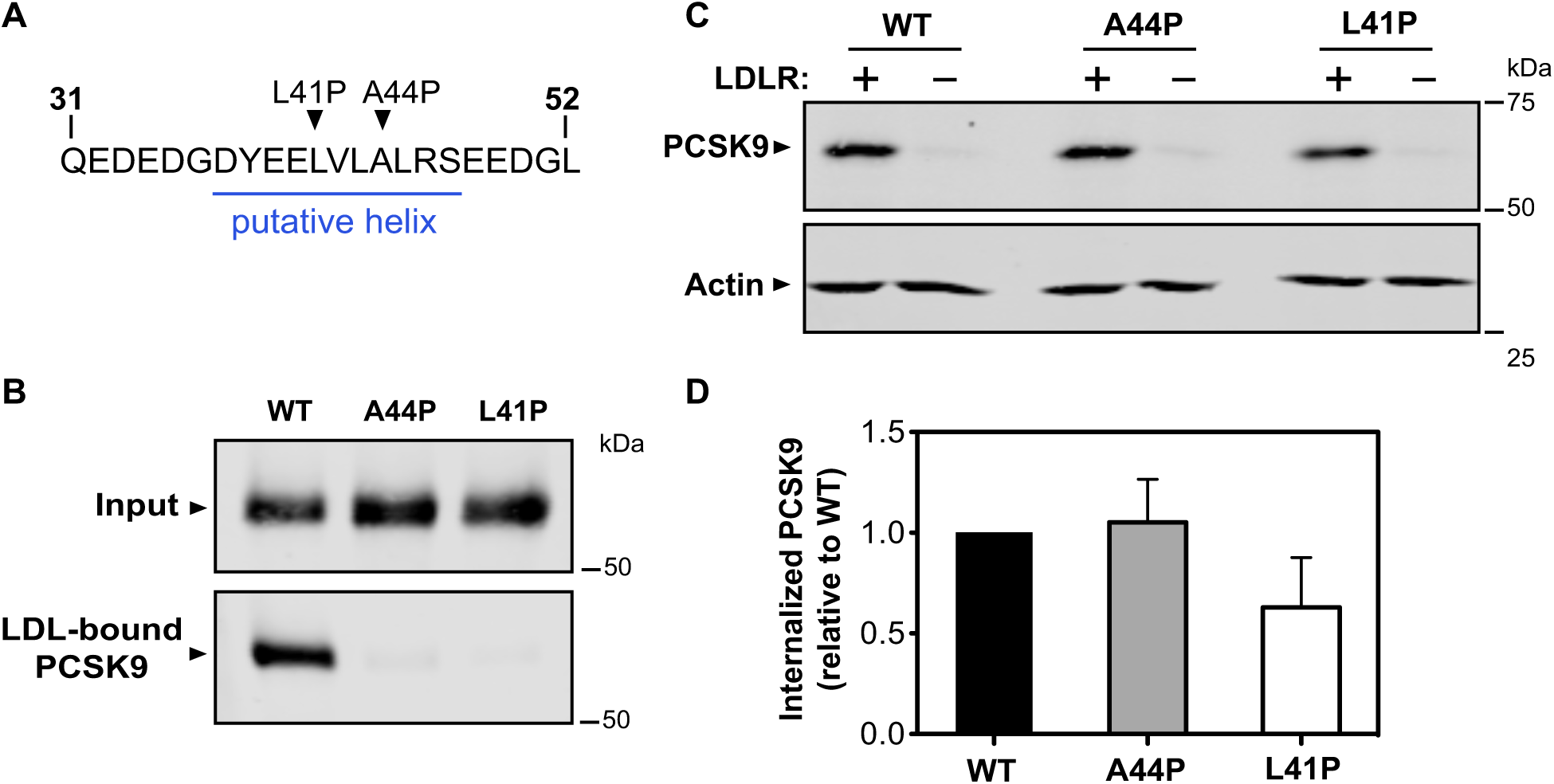
Proline substitutions in PCSK9 N-terminus prevent LDL, but not LDLR, binding. (**A**) Position of proline substitutions in putative helical segment. (**B**) Analysis of *in vitro* PCSK9-LDL binding reactions. Conditioned cell culture medium containing wild-type PCSK9 (WT) or proline mutants (A44P or L41P) were incubated with LDL prior to density gradient-ultracentrifugation and Western blot analysis of LDL-containing fractions. (**C**) HEK293 cells were transfected with LDLR (+) or vector control (-) and incubated with conditioned medium containing PCSK9 proteins. Cell uptake of PCSK9 was measured by western blotting. (**D**) Densitometric analyses of Western blot in *C*. Error bars represent SEM (n=3).

### Helical conformation specifically affects PCSK9 binding affinity to LDL

To more quantitatively assess the effect of the A44P mutation in full-length PCSK9, we performed a steady-state binding analysis in which fluorescent dye-labeled WT-PCSK9 was incubated with LDL in the presence of increasing concentrations of unlabeled WT-PCSK9 or PCSK9-A44P. For comparison purposes, we also included PCSK9 lacking the N-terminal 21 residues (Δ31-52), which is defective in LDL binding (18) and hyperactive in terms of LDLR binding/degradation (26,34). Competition binding curves showed that the PCSK9-A44P mutant had a >5-fold decrease in LDL binding affinity compared to WT-PCSK9 (Figures 5A and B), whereas Δ31-52–PCSK9 affinity for LDL was more than 2 orders of magnitude lower than that of WT-PCSK9 (Figure 5A). We also assessed the affinity of PCSK9-A44P for the purified LDLR extracellular domain (LDLR-ECD), which was decreased ∼20% compared to WT-PCSK9 (Figure 5C). Nevertheless, when added to the medium of cultured HepG2 cells, PCSK9-A44P was able to mediate dose-dependent degradation of cell surface LDLRs at levels comparable to WT-PCSK9 (Figure 5D), in agreement with a recent study (28). Collectively, these data support that helical conformation in the prodomain N-terminus plays a larger role in LDL binding relative to its role in PCSK9-LDLR association and LDLR degradation.

**Figure 5.**
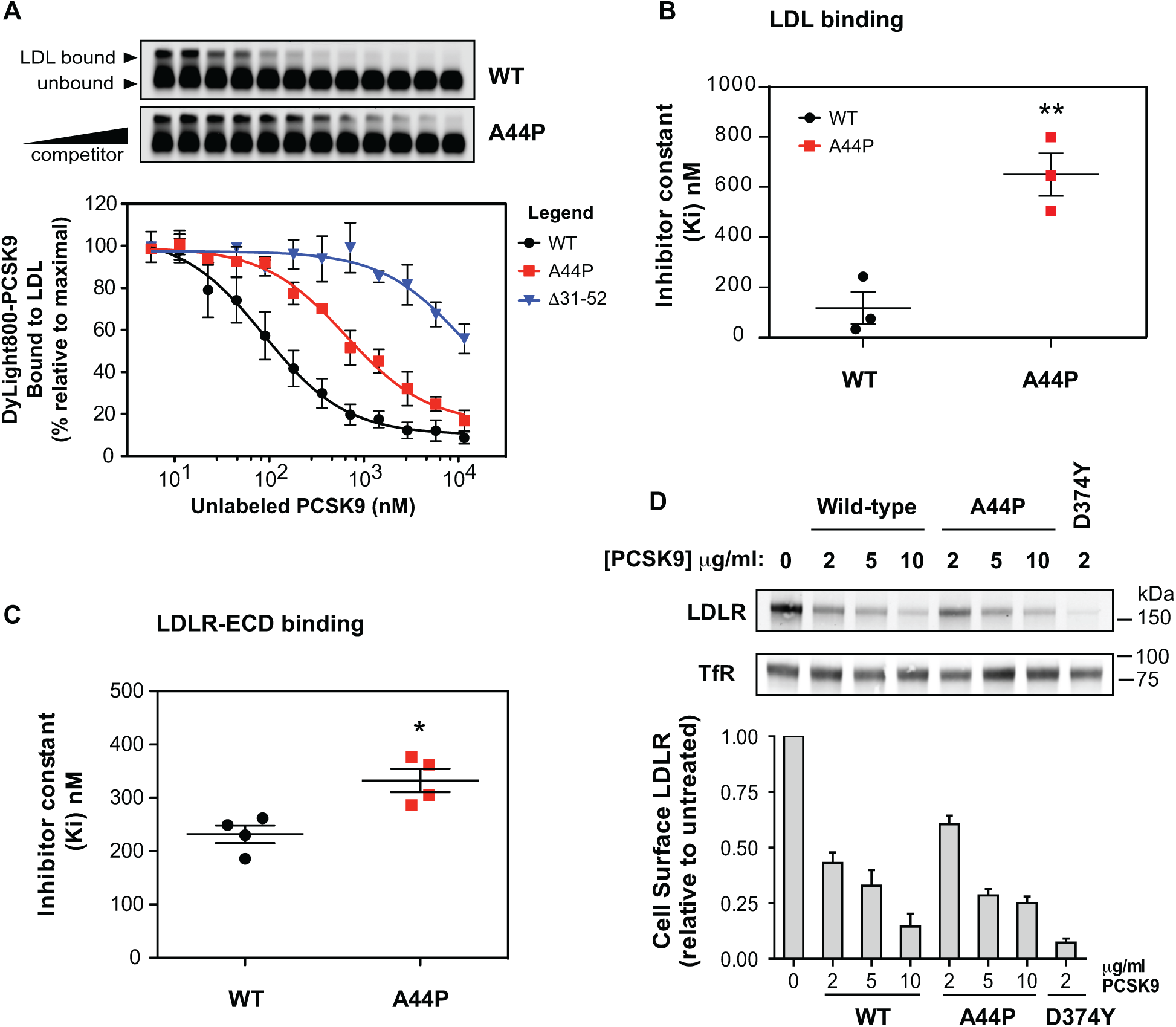
Helix-breaking A44P mutation in PCSK9 selectively inhibits LDL binding. (**A**) In vitro competition binding of WT, A44P and Δ31-52 forms of PCSK9 to LDL. LDL particles were incubated with Dylight800-labeled PCSK9 in the presence of increasing concentrations of unlabeled competitor proteins. Reaction mixtures were separated on agarose gels (*top*) and fluorophore-labeled PCSK9 binding to LDL was quantified and fitted to competition binding curves using non-linear regression (*bottom*). (**B**) Inhibitor constants (Ki) obtained from curves in *A*. Error bars represent SEM (n=3). Significant change in LDL binding of A44P mutant PCSK9 compared with WT-PCSK9 was determined by Student’s t-test: **, *p* < 0.01. (**C**) Competition binding of WT-PCSK9 and A44P mutant to LDLR extracellular domain (LDLR-ECD). Shown is graphical representation of inhibitor constants (Ki) obtained from binding curves. Error bars represent SEM (n=4). Significant change in LDLR-ECD binding of A44P mutant PCSK9 compared with WT-PCSK9 was determined by Student’s t-test: *, *p* < 0.05. (**D**) Dose response of cell surface LDLR degradation in HepG2 cells cultured 4 h in medium containing increasing concentrations of WT or A44P forms of PCSK9. Biotinylated cell surface LDLRs were isolated and quantified by western blotting (*top*) using transferrin receptor (TfR) as a loading control. Densitometric data (*bottom*) was compared to untreated cells. LDLR degradation by GOF mutant D374Y PCSK9 was used for comparison and as a positive control. Error bars represent SEM (n=3).

### R46L substitution increases helix propensity in prodomain-derived peptides

The natural PCSK9 mutation R46L is a LOF variant associated with lowered plasma PCSK9 levels, an anti-atherogenic lipid profile and decreased incidence of CHD (5,35,36). Notably, an R46L substitution would extend the nonpolar face of the predicted AH motif in the IDR and reorient and increase the amplitude of the hydrophobic moment, as shown in helical wheel models (Figure 6A). Helical net projections show the R46L mutation is also predicted to add additional side-chain interactions to stabilize the helical conformation (Figure 6B). In agreement with these models, structural analysis using CD spectroscopy showed that an R46L-containing peptide representing residues 36-47 in PCSK9 presented a much stronger helical shift in a DPC micelle environment, with changes to both the intensity and position of the minimum relative to the corresponding WT peptide (Figure 6C). The percent helicity of the peptides obtained at different DPC micelle concentrations allowed micelle binding curves to be obtained, revealing that the R46L substitution conferred an ∼8-fold higher affinity for micelles compared to the WT peptide (Figures 6D and E). The R46L mutation in full-length PCSK9 did not affect binding affinity to isolated LDL particles, as unlabeled versions of WT-PCSK9 and PCSK9-R46L showed near equal abilities to compete with dye-labeled PCSK9 for binding to LDL (Supplemental Figure S2).

**Figure 6.**
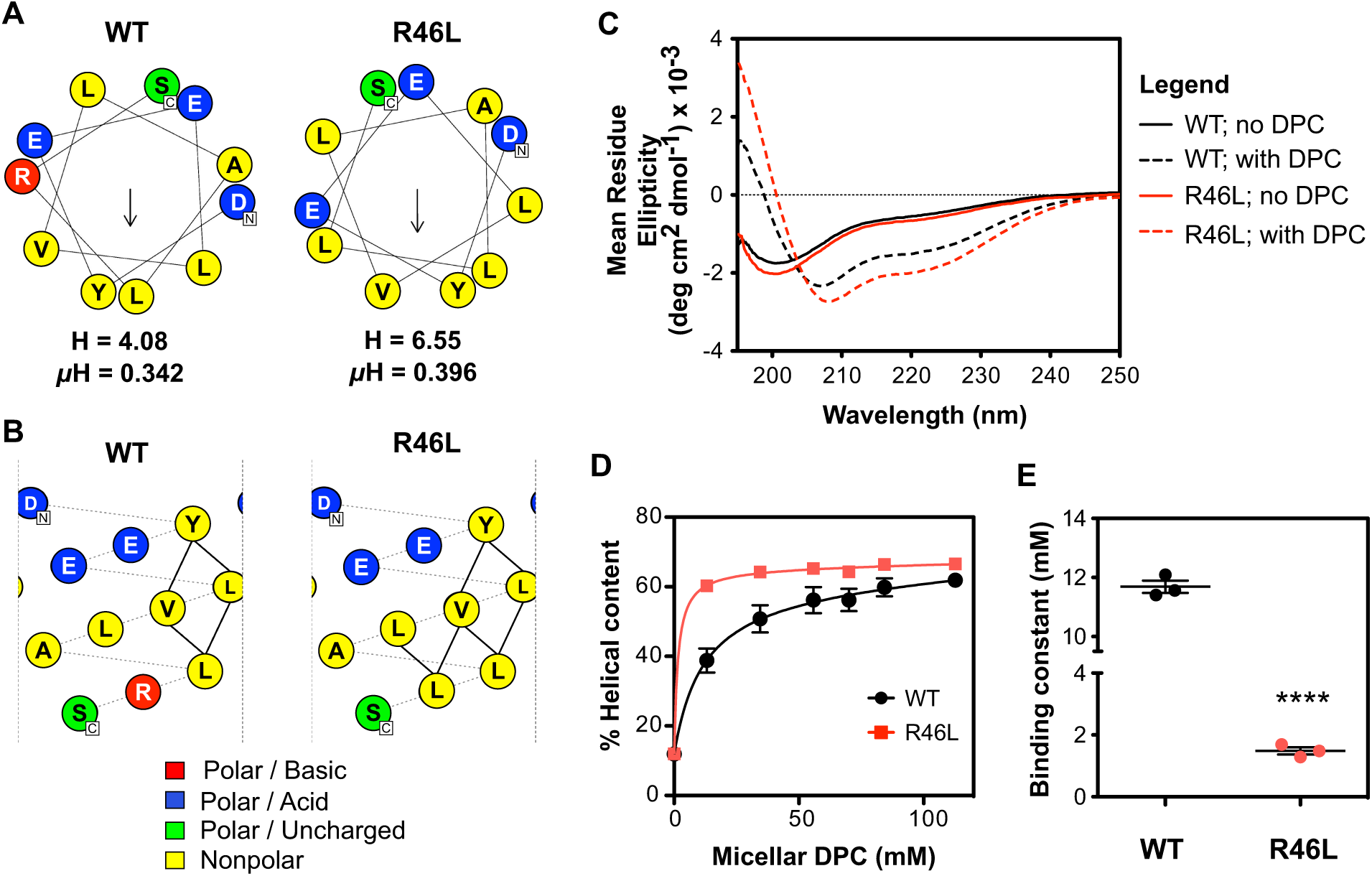
Natural mutation R46L modulates helicity in the prodomain N-terminal region. **(A)** Helical wheel representation of amphipathic α-helices (residues 37-47) generated from HeliQuest with either arginine or leucine at position 46. The *arrow* indicates the magnitude and direction of the hydrophobic moment (32). The hydrophobicity (H) and hydrophobic moment (μH) from HeliQuest are also shown. **(B)** Predicted hydrophobic side-chain interactions (*solid lines*) between residues of the putative N-terminal helix with either arginine or leucine at position 46. Constructed on NetWheels software online (http://lbqp.unb.br/NetWheels/). **(C)** Circular dichroism spectra prodomain peptides either wild-type: GDYEELVLALRS (*black*), or R46L: GDYEELVLALLS (*red*) in the absence (*solid lines*) or presence (*dashed lines*) of DPC micelles (n=3). **(D)** Mean % helical content of either the wild-type or the R46L variant peptide as DPC concentration is increased. Secondary structure calculations based on circular dichroism spectra using deconvolution algorithm CONTIN. Error bars represent SEM (n=3). **(E)** Binding constants obtained from curves in *D*. Error bars represent SEM (n=3). Significant change in micelle binding compared with WT peptide was determined by Student’s t-test: ****, *p* < 0.0001.

### Tyr-38 sulfation does not prevent PCSK9-LDL association

Numerous plasma proteins, such as exchangeable apolipoproteins, employ AH motifs for reversible binding to circulating lipoproteins (37). This function involves direct membrane association in which the nonpolar face of the helix intercalates between the fatty acyl chains in the single-layer phospholipid outer membrane of the lipoprotein. The transient AH in the PCSK9 prodomain has a comparatively small nonpolar face consisting of four residues, including Tyr-38 identified as a site of O-sulfation (38). We took advantage of this post-translational modification to test if introduction of a negative sulfate group to the nonpolar face in the helix affected PCSK9’s ability to bind LDL. HEK293 cells transiently over-expressing various PCSK9 constructs were incubated with Na-[^35^S]O_4_ and radiolabeled PCSK9 was immunoprecipitated for SDS-PAGE analysis. As previously reported (38), a PCSK9-Y38F was only [^35^S]-labeled at a site in the catalytic/CHR domain, confirming Tyr-38 as the sole sulfation site in the prodomain under these experimental conditions (Supplemental Figure S3A). Following incubation with LDL, we were able to readily detect LDL-bound PCSK9 containing sulfated Tyr-38, whereas PCSK9-A44P containing sulfated Tyr-38 was defective in LDL binding (Supplemental Figure S3B). This result further supports that helical structure in the prodomain N-terminus is essential for LDL binding and also indicates that an uninterrupted hydrophobic face in the helix is not an absolute requirement. Nevertheless, hydrophobicity at this site could assist in formation of an AH motif in PCSK9 that facilitates LDL binding. In support, hydrophobic substitutions (Y38L or Y38F) for the Tyr-38 residue were found to largely preserve LDL-binding while lower hydrophobicity (Y38A) caused a significant decrease of >50% (Supplemental Figure S3C).

### Evidence of an interdomain interaction required for PCSK9-LDL binding

The inability of Tyr-38 sulfation to prevent LDL binding (Supplemental Figure S3B) suggests that formation of the transient AH motif in the N-terminal region of PCSK9’s prodomain promotes LDL association, although not necessarily through the membrane insertion of its hydrophobic face. Alternatively, the helical conformation could facilitate a protein-protein interaction, either directly with LDL-apoB100 or to an intramolecular site. To assess the latter possibility, computational molecular modeling of PCSK9 to include the missing IDR in known structures was performed using the ab initio structure prediction method implemented in ROSETTA (39,40). Constraints from the highest-resolution PCSK9 x-ray crystal structure (PDB ID 2QTW) (29) were applied to conform to known parts of the protein, whereas an exhaustive conformational sampling was carried out to model residues 31 to 60. The resultant model in Figure 7A shows the N-terminal prodomain inserted into a central cavity formed by the prodomain, the catalytic domain and the CHR domain, stabilized in this position by an intramolecular interaction between an extended N-terminal helix (aa 32-48) and the CHR domain. The model also predicts adoption of helical conformation in a second sequence (aa 50-58) adjacent to a Ser-47 phosphorylation site (41) and within close proximity to a β-strand of the prodomain (denoted β_0_) described as unique in PCSK9 compared with all known structures of subtilisn-like enzymes (11). The Arg-46 residue is located in an otherwise hydrophobic region of the N-terminal helix and does not interact with the rest of the protein; thus, it would be expected that the R46L LOF mutation in PCSK9 would expand a hydrophobic face of the helix. The intramolecular interaction predicted by our model involved negative-charged residues (Glu-32, Asp-35 and Glu-39) and aromatic Tyr-38 in the helical N-terminal prodomain conformation and multiple positive-charged residues in the CHR domain (Arg-496, Arg-499, Arg-510, His-512 and His-565). Several GOF PCSK9 mutations associated with FH (R469W, R496W and F515L) are clustered on a surface region of the CHR domain at or near the predicted interdomain interface (Figure 7A, inset). When introduced into recombinant human PCSK9, all three mutations greatly inhibited LDL binding in vitro (Figures 7B and C). In particular, recombinant PCSK9-R496W appeared to be completely unable to bind LDL, whereas R469W and F515L substitutions lowered binding by ∼60-70% (Figure 7C). None of these mutations greatly affected PCSK9 secretion into cell culture medium (Figure 7B), in agreement with a recent study (42), and all exhibited normal binding to cell surface LDLRs overexpressed in HEK293 cells (Figure 7D).

**Figure 7.**
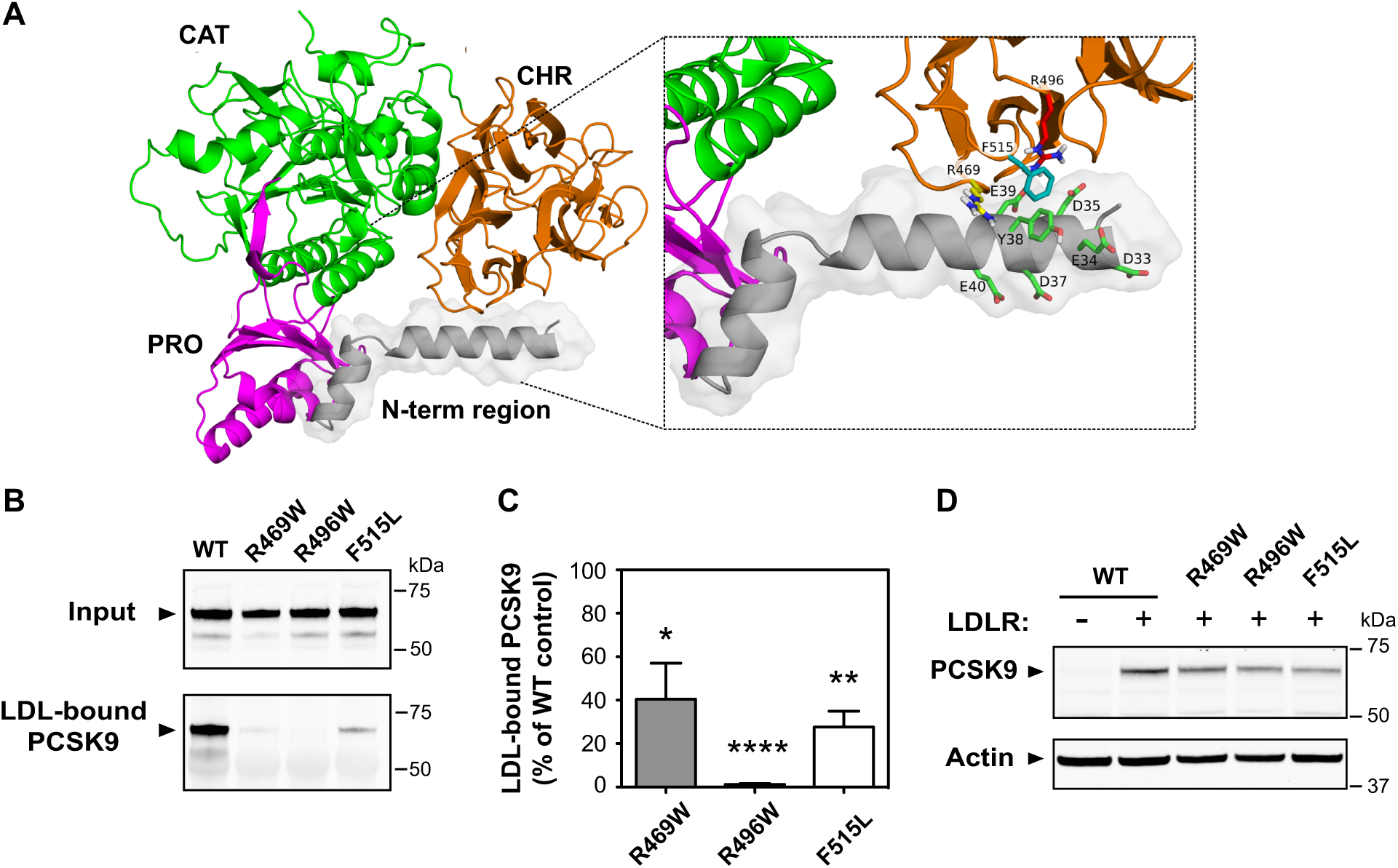
GOF *PCSK9* mutations in the C-terminal domain disrupt LDL binding *in vitro*. **(A)** Structure of full-length PCSK9 modeled by ROSETTA with constraints to conform to known parts of the structure from the highest-resolution x-ray crystal structure (PDB ID 2QTW) (29). Color scheme: Prodomain (Pro, magenta), catalytic domain (Cat, green), cysteine-histidine rich (CHR) domain (CHR, orange), and N-terminal region of the prodomain (grey with transparent surfaces). Box: details of prodomain/CHR domain interface; side-chain sticks (green) highlight polar residues in the N-terminal region; side-chain sticks for R469 (yellow), R496 (red) and F515 (cyan) show the close proximity of residues associated with ADH. **(B)** Analysis of *in vitro* PCSK9-LDL binding reactions. Conditioned medium containing wild-type PCSK9 (WT) or GOF variants R469W, R496W and F515L were incubated with LDL prior to density gradient-ultracentrifugation to isolate an LDL fraction and visualization of PCSK9 by western blot. (**C**) Densitometric analyses of western blot in *C.* Error bars represent SEM (n=4). Significant change in LDL binding compared to WT-PCSK9 control (set to 100%) was determined by One sample t-test: * *p* < 0.05; ** *p* < 0.01; **** *p* < 0.0001. (**D**) HEK293 cells were transfected with LDLR (+) or vector control (-) and incubated with conditioned medium containing the indicated PCSK9 proteins. Cell uptake of PCSK9 was visualized by western blotting. Shown is a representative experiment (n=3).

## DISCUSSION

IDRs in proteins frequently have crucial roles in regulatory and signaling processes due to increased flexibility and accessibility for post-translational modification and protein-protein interactions, and in some cases undergo disorder-to-order structural transition as part of this function (43-45). We previously reported that association of PCSK9 with LDL particles *in vitro* is dependent on an IDR in the prodomain N-terminus (18). In the current study, we provide biochemical, structural and genetic evidence that this requirement is manifested as a transient AH, as follows: 1) an N-terminal prodomain-derived peptide adopted α-helical conformation in a membrane-mimetic environment (DPC micelles) (Figure 3A), which was greatly enhanced by an R46L substitution representing an athero-protective *PCSK9* LOF mutation (Figure 6); 2) a helix-breaking proline substitution (A44P) in PCSK9 lowered LDL binding affinity by >5-fold (Figures 5A and B); 3) a computational 3D model of PCSK9 predicted that α-helical structure in prodomain N-terminus aligns multiple polar residues to interact with the CHR domain (Figure 7A); and 4) several *PCSK9* GOF mutations located at or near the predicted interface in the CHR domain greatly diminished LDL binding (Figure 7B). Considered together, we propose a two-step process by which allosteric conformational changes in PCSK9 facilitate LDL binding: *i*) the initial encounter with LDL in circulation provides hydrophobic lipid and/or protein contacts to trigger AH formation in the N-terminal IDR; *ii*) this structural transition promotes an interdomain interaction involving the prodomain and CHR domain which in turn allows stable LDL association.

GOF mutations in *PCSK9* are rare in human populations and can result in substantially increased plasma LDL-C levels in FH patients (46). As such, delineation of underlying mechanisms can provide clear insight on aspects of PCSK9 function (6). In Figure 7B we identify several GOF mutations in PCSK9 that severely inhibit (R469W and F515L) or abolish (R496W) LDL binding *in vitro*. The affected amino acids are clustered on a surface region in CHR domain module (CM)1 (aa 453– 531), one of three structurally similar subdomains arranged in a pseudothreefold axis (29). This same site in CM1 was previously shown to engage a monoclonal antibody that inhibited PCSK9-mediated LDLR degradation in cells and *in vivo* without affecting binding affinity to the LDLR EGF-A domain (47). Although the precise mechanism of PCSK9-LDL binding remains to be determined, Arg-496 could be a key residue in a prodomain-CM1 intramolecular interaction that serves as an essential component. In support, computational modeling studies predicted that adoption of helical conformation in the prodomain IDR aligns potential interactions between Asp-35, Tyr-38 and Glu-39 with Arg-496 in CM1 (Figure 7A). The Arg-469 and Phe-515 residues do not interact with the prodomain in the model and possibly are more directly involved in binding to the LDL particle.

The R496W *PCSK9* mutation was first identified in an individual also heterozygous for an *LDLR* LOF mutation (E228K), the combined effect being severe hypercholesterolemia and early-onset cardiovascular heart disease (48). A high frequency of R496W *PCSK9* mutations (∼9%) was recently reported in a Turkish FH cohort, associated with highly elevated plasma LDL-C levels and early onset CHD in 7 of 8 cases (49). In a multi-center study of FH associated with various *PCSK9* GOF mutations, the R496W mutation was among the more severe in terms of elevated LDL-C levels (46). Like R496W, the R469W and F515L *PCSK9* mutations have been identified in FH patients (35,50), but functional effects have remained unknown. In cell culture studies, all three GOF mutations exhibited only minor effects on maturation/secretion of PCSK9 and on LDLR degrading ability (42,51,52). In the current study, binding of PCSK9 to LDLRs overexpressed in HEK293 cells was largely unaffected (Figure 7D). Given the lack of other functional effects, it is highly probable that defective LDL binding underlies the GOF phenotype associated with these *PCSK9* mutations in the CHR domain. Since PCSK9 activity raises plasma LDL levels, inhibition via direct binding to LDL particles could provide a form of negative feedback control. In this scenario, PCSK9 variants that bind poorly to LDL would be less subject to this inhibition, resulting in FH.

Lipid-ordered AHs are archetypal membrane-binding motifs found in numerous plasma proteins that, like PCSK9, reversibly associate with circulating lipoproteins and regulate lipoprotein metabolism (33,37). However, the current data supports that the primary function of a transient AH in PCSK9 is not membrane insertion, but rather to align an interdomain interaction that in turn facilitates LDL binding. This role in a protein-protein interaction is more akin to transient AH motifs found in acidic activation domains (AADs) of transcription factors. Most AADs are intrinsically disordered, which facilitates association with multiple partner proteins in an amorphous and flexible manner (53). In some cases, coupled folding and binding occurs driven by weak hydrophobic “encounter” interactions that initiate helix formation (54). A paradigm for the evolution of transient helical motifs in signaling proteins of higher organisms is the tumour suppressor p53, in which formation of short AH interaction motifs in its AAD modulates target protein binding and transcriptional control (55-57). In a similar manner, order/disorder transition in the prodomain IDR of PCSK9 could affect the stringency or affinity of its interactions with potential binding partners, including other intramolecular sites.

A previous study showed that PCSK9 lacking an N-terminal portion of the prodomain IDR (Δ31-52) bound LDLR *in vitro* with >7-fold higher affinity, identifying a negative allosteric function (26). Further studies narrowed this effect to a highly acidic segment (aa 32-40; EDEDGDYEE) hypothesized to participate in intramolecular electrostatic interactions in a cryptic autoinhibited conformation, perhaps involving the nearby CHR domain (27,30,58). Indeed, a predicted inter-domain interface site in CM1 (Figure 7A) contains a cluster of arginine residues at positions 469, 496, 499 and 510 that could interact with acidic amino acids in a disordered prodomain N-terminus. This may be sufficient to maintain PCSK9 in an autoinhibited state under basal conditions, explaining why a helix-breaking A44P mutation in PCSK9 did not increase LDLR binding/degradative functions (Figures 5C and D). On the other hand, adoption of rigid α-helical structure could favor certain contacts, possibly involving the Arg-496 residue identified as key to LDL binding (Figure 7B). If so, this would explain why PCSK9-A44P had lowered LDL binding affinity but was not as defective in this function as Δ31-52–PCSK9, in which an interaction with CM1 would be completely abolished (Figures 5A and B). Indeed, LDL association may stabilize an autoinhibitory prodomain-CM1 interaction in PCSK9 that otherwise can be released by activating factors. Two such factors are known to improve the LDLR binding function of PCSK9, namely acidic pH (11,16,22) and its association with heparin-sulfate proteoglycans (HSPGs) at the surface of hepatocytes (59). HSPGs were recently shown to modulate the ability of LDL to inhibit PCSK9 activity, suggesting an interplay between these regulatory factors (23).

The N-terminal prodomain IDR is the site of several *PCSK9* mutations associated with plasma LDL-C levels in humans (6,27) including an athero-protective R46L *PCSK9* allele found in ∼3% of Caucasians (5,36,60). Unlike more deleterious LOF mutations that disrupt autocatalytic cleavage or protein folding, R46L does not affect PCSK9 secretion (61) and only minor effects of this mutation on cellular LDLR binding and degradation have been reported (22,58,62). As shown in Figure 6, this mutation is predicted to extend the hydrophobic face of a transient AH motif, and an R46L substitution in a prodomain-derived peptide had a substantial positive effect on coil-to-helix structural transition in the presence of DPC micelles, a membrane-mimetic. Thus, increased helix propensity could contribute to the LOF phenotype, perhaps via enhanced plasma clearance of PCSK9-R46L as a passive component of LDL particles. In support, individuals carrying the R46L allele have significantly lowered plasma PCSK9 concentrations (35). Counter to this hypothesis, we found that purified recombinant PCSK9-R46L displayed normal LDL binding affinity *in vitro* (Supplemental Figure S2). However, However, helix formation in the WT-PCSK9 prodomain may not be limiting under these in vitro conditions, masking any differences. It should also be noted that the *PCSK9* R46L allele in humans is associated with only modest lowering of plasma LDL-C (∼15%) with athero-protection likely due to life-long exposure (5,60). Thus, any acute functional effects of R46L in full-length PCSK9 are likely to be minor, highlighting the utility of structural approaches for mechanistic insight.

The prodomain IDR is also the site of two post-translational modifications, namely phosphorylation at Ser-47 (41) and sulfation at Tyr-38 (38). Although Tyr-38 sulfation introduces a negative charge within the hydrophobic face of the transient AH motif (Figure 2D), it did not affect LDL binding *in vitro* (Supplemental Figure S3B). On the other hand, mutation of Tyr-38 to alanine lowered PCSK9-LDL binding by >50%, whereas substitutions of higher hydrophobicity (Phe or Leu) preserved LDL binding (Supplemental Figure S3C). A possible explanation is that decreased hydrophobicity upon Tyr-38 sulfation is compensated by direct involvement of the sulfo-tyrosine residue in the prodomain-CM1 interaction that controls LDL binding. In a recent study, a monoclonal antibody was shown to specifically bind and stabilize helical conformation in peptides derived from the N-terminal IDR in the PCSK9 prodomain (28). Binding affinity was greatly enhanced by sulfation of Tyr-38 (28), consistent with a known role of sulfo-tyrosine in promoting protein-protein interactions (63,64). Post-translational modifications often modulate the functional interactions of short interaction motifs within IDRs in proteins (65); thus, Tyr-38 sulfation could serve this role in the N-terminal prodomain IDR in PCSK9.

Determining the functional outcome of PCSK9-LDL association is crucial to our overall understanding of reciprocal feedback mechanisms that control plasma concentrations of LDL-C and associated risk of cardiovascular heart disease. It is established that LDLR-mediated uptake and degradation of LDL particles in hepatocytes raises intracellular levels of cholesterol and suppresses activity of the SREBP-2 transcription factor, thereby lowering gene expression of both LDLR and PCSK9 (3,66). Also, LDLR-mediated uptake of PCSK9 results in LDLR degradation and reduced LDL and PCSK9 clearance from plasma (13). The data in the current study supports the existence of an additional feedback loop whereby circulating PCSK9 activity is inhibited via its direct interaction with LDL particles in response to intravascular LDL accumulation. The identification of point-mutant forms of PCSK9 selectively defective for LDL association, such as A44P, provides important molecular tools to further test this hypothesis in animal models of dyslipidemia.

## EXPERIMENTAL PROCEDURES

### Materials

We obtained fetal bovine serum (FBS) and newborn calf serum from ThermoFisher, Optiprep™ density gradient medium (60% w/v iodixanol) from Axis-Shield, EDTA-free Complete™ Protease Inhibitor Tablets were from Roche, NP-40 detergent was from Biovision and PolyJet DNA transfection reagent was from FroggaBio (Toronto, Ontario, Canada). All other reagents were from Sigma-Aldrich unless otherwise specified. Sodium mevalonate was prepared from mevalonic acid as described (67). Newborn calf lipoprotein-deficient serum (NCLPDS) (d > 1.215 g/ml) was prepared by ultracentrifugation (68). The LDLR cDNA expression vector used in these studies was pLDLR17 (69).

### Cell culture

HEK293 and HepG2 cells (American Type Culture Collection) were maintained in monolayer culture at 37°C and 5% CO_2_ in one of the following medium: Medium A contained DMEM (4.5 g/L glucose; Gibco) supplemented 100 U/ml penicillin and 100 µg/ml streptomycin sulfate; Medium B contained Medium A supplemented with 10% FBS (v/v); Medium C contained Medium A supplemented with 5% (v/v) NCLPDS; Medium D contained DMEM (1.0 g/L glucose; Gibco) supplemented 100 U/ml penicillin and 100 µg/ml streptomycin sulfate; sterol-depleting Medium E contained Medium D supplemented with 5% (v/v) NCLPDS, 10 µM pravastatin, and 50 µM sodium mevalonate.

### Protein purification and labeling

FLAG epitope-tagged recombinant human wild-type PCSK9 along with A44P, R46L and Δ31-52 variants were produced in stably-transfected HEK293S cells and purified from conditioned culture medium as previously described (18). Fluorescently labeled wild-type PCSK9 was prepared using the DyLight800 Antibody Labeling Kit (ThermoFisher) as per manufacturer’s instructions followed by gel filtration chromatography on a Superdex 200 10/300 GL column (GE Healthcare) to remove unbound dye. Extracellular domain (ECD) of LDLR used for PCSK9 binding assays was partially purified from conditioned medium of HEK293S cells cultured as described (18) stably transfected with a plasmid encoding amino acids 1-692 of human LDLR containing a C-terminal 6X-His tag (a kind gift from R. Milne, U. of Ottawa). LDLR-ECD was enriched by affinity chromatography using TALON superflow affinity resin (Clontech) followed by gel filtration chromatography on a Superdex 200 10/300 GL column (GE Healthcare).

### Antibodies

The following antibodies were used for western blotting: A mouse hybridoma clone expressing monoclonal antibody 15A6 recognizing an epitope in the C-terminal CHR domain of PCSK9 was a generous gift from J. Horton (UTSouthwestern Medical Center, Dallas, TX). The IgG fraction was purified from hybridoma culture medium by Protein A chromatography on a Profinia™ protein purification system (Bio-Rad) as per manufacturer’s instructions. A rabbit anti-serum 3143 against the C-terminal 14 amino acids of LDLR was the kind gift of J. Herz (UTSouthwestern Medical Center). Anti-actin mouse monoclonal ascites fluid (clone AC-40) was from Sigma. Mouse anti-human transferrin receptor antibody was from Life Technologies. Secondary IRDye-labeled goat anti-mouse and anti-rabbit IgG antibodies were from LI-COR Biosciences. Rabbit anti-serum 1697 raised against full-length human PCSK9 was used for immunoprecipitation and was custom produced by Biomatik (Cambridge, Ontario, Canada)

### Western blotting

Samples were loaded onto 8% SDS-polyacrylamide gels or 4–12% Tris/HEPES-SDS Bolt™ precast gels (Thermo-Invitrogen) for electrophoresis. The size-separated proteins were then transferred to nitrocellulose membranes (Bio-Rad). Primary antibodies (described above) and IRdye800-conjugated secondary antibodies were used to detect target proteins. Signal was detected using the LI-COR Odyssey infrared imaging system (LI-COR Biosciences). Quantification of the intensity of the bands was obtained using Odyssey 2.0 software (LI-COR Biosciences).

### Site-directed Mutagenesis

The pcDNA3-PCSK9-FLAG vector (70) codes for full-length human wild type PCSK9 with a FLAG-tag epitope attached at the C-terminus. This vector was used as the template to generate PCSK9 mutants. Mutagenesis was carried out using the QuikChange site-directed mutagenesis protocol from Stratagene (La Jolla, CA). PCR primers were custom-synthesized by Invitrogen-Life Technologies or Integrated DNA Technologies. The primers used for generation of the mutants are listed in Supplemental Table 1. All desired mutations and absence of extraneous mutations were confirmed by sequencing the entire coding region.

### Preparation of Conditioned Media

HEK293 cells cultured in Medium B to ∼70% confluency in a 100-mm Petri dish were transfected with a total of 3 μg of PCSK9 cDNA expression vectors using PolyJet DNA transfection reagent as per manufacturer’s instructions. At 18 h post-transfection, the cells were washed and incubated with serum-free Medium A containing 1X ITS (insulin-transferrin-selenium) cell culture supplement. Conditioned media were recovered 48 h post-transfection and buffer exchanged with HBS-C (HEPES buffered saline-calcium buffer: 25 mM HEPES-KOH, pH 7.4, 150 mM NaCl, 2 mM CaCl_2_) and concentrated ∼10-fold on an Amicon Ultra-4 centrifugal filter unit with a 10-kDa membrane cutoff (Millipore). PCSK9 in conditioned media preparations was quantified by western blot analysis using purified PCSK9 as a standard and stored in single-use aliquots at −80°C.

### LDL isolation

All procedures using human subjects received regulatory approval from the Human Research Ethics board at the University of Ottawa Heart Institute. Blood samples were drawn from fasted healthy volunteers into evacuated tubes containing EDTA and plasma was separated by low speed centrifugation. Protease inhibitors (1 mM PMSF, 50 U/ml aprotinin and Complete™ protease inhibitor cocktail) and antioxidant (20 μM butylated hydroxytoluene) were added to the cleared plasma. LDL (d = 1.019-1.065 g/ml) was isolated using sequential potassium bromide flotation ultracentrifugation (71) followed by extensive dialysis against phosphate-buffered saline (PBS) containing 0.25 mM EDTA. LDL was stored at 4°C protected from light and used within 2-3 weeks. Alternatively, LDL was stored at −80°C in 10% (w/v) sucrose as described (72), which did not affect subsequent PCSK9 binding tests.

### PCSK9-LDL binding assay

Binding reactions (1.0 ml volume) each containing 500 μg LDL and approximately 1 µg PCSK9 (in concentrated conditioned media) and 0.5% BSA (w/v) in HBS-C buffer (25 mM HEPES-KOH, pH 7.4, 150 mM NaCl, 2 mM CaCl_2_) were incubated at 37°C for 1 hour. LDL-bound and free PCSK9 were then separated by Optiprep gradient ultracentrifugation according to a modified version of a previously described protocol (73). Briefly, a 9% Optiprep sample solution was prepared by diluting 1.0 ml of each binding reaction with 0.45 ml of 60% Optiprep and 1.55 ml of 25 mM HEPES-KOH, pH 7.4. The 9% sample was overlayed with 25mM HEPES-KOH, pH 7.4 in a 3.3 ml Optiseal tube (Beckman). Tubes were centrifuged in a TLN100 rotor (Beckman Coulter) at 100,000 rpm for 2 hours at 4°C. LDL containing fractions (600 μl) were collected by tube puncture into a 1 ml syringe. A one-fifth amount of input (original binding reaction before Optiprep addition) or the LDL-fraction were immunoprecipitated for 18 h at 4°C using anti-PCSK9 rabbit polyclonal 1697 (see above) and anti-rabbit Trueblot agarose beads (Rockland). The collected beads were washed and immunoprecipitated proteins were resolved by SDS-PAGE on 8% acrylamide gels. PCSK9 content was then quantified by western blotting and values were normalized to reactions containing wild-type PCSK9.

### Competition binding curves

For PCSK9-LDL binding analysis, 0.5 mg/ml of LDL was incubated with 10 nM of DyLight800-labeled wild-type PCSK9 in HBS-C buffer containing 1% (w/v) BSA for 90 min at 37°C with increasing amounts of unlabeled wild-type or mutant PCSK9. 4X loading dye (10% Ficoll-400, 0.01% bromophenol blue) was added to the reactions and the mixtures were resolved on 0.7% agarose gels (SeaKem LE) for 2h at 40V with electrode buffer of 90 mM Tris, pH 8.0, 80 mM borate, 2 mM calcium lactate. Gels were scanned directly using the LI-COR Odyssey infrared imaging system (LI-COR Biosciences) and bands representing DyLight800-PCSK9 bound to LDL were quantified using Odyssey 2.0 software (LI-COR Biosciences). For PCSK9-LDLR binding analysis, partially purified LDLR-ECD (see above) was diluted in TBS-C (50 mM Tris pH 7.4, 90 mM NaCl, 2 mM CaCl_2_) and blotted directly onto 0.45 μm pore size nitrocellulose membranes (Bio-Rad) using a BioDot SP slot-blot apparatus (BioRad) according to manufacturer’s instructions and blots were blocked for 30 min in TBS-C containing 5% (w/v) non-fat milk. All incubations (90 min) and subsequent washes (3 x 15 min) were carried out at room temperature with gentle oscillation in TBS-C buffer containing 2.5% (w/v) non-fat milk. Blots were incubated with 0.1 μg/ml DyLight800-labeled wild-type PCSK9 with increasing amounts of unlabeled wild-type or mutant PCSK9. Blots were then washed and scanned using the LI-COR Odyssey Infrared Imaging System and band intensity was quantified using Odyssey v2.0 software (LI-COR Biosciences). The amount of competitor PCSK9 protein required for 50% inhibition of fluorophore-labeled PCSK9 binding (Ki) was determined by fitting data to a sigmoidal dose response curve using nonlinear regression (GraphPad Prism 5 software).

### PCSK9 Cellular uptake assays

HEK293 cells were cultured in Medium B in 6-well dishes to 70% confluency, then transiently transfected with 1 μg wild-type human LDLR plasmid per well using PolyJet DNA transfection reagent as per manufacturer’s instructions. The following day, cells were switched to lipoprotein-deficient Medium C and treated for 2 hours at 37°C with 10 μg/ml PCSK9 (from conditioned media). Cells were then washed in ice-cold PBS-CM (PBS with 1 mM MgCl2 and 0.1 mM CaCl2) and whole cell extracts were prepared in Tris Lysis Buffer (50 mm Tris-Cl, pH 7.4, 150 mm NaCl, 1% NP-40, 0.5% sodium deoxycholate, 5 mm EDTA, 5 mm EGTA, 1X Complete™ protease inhibitor cocktail, 1 mM phenylmethylsulfonyl fluoride). Extracted proteins were resolved on 8% SDS-PAGE and LDLR, PCSK9 and actin were detected by western blotting.

### Cell surface LDLR degradation assays

HepG2 cells in 60 mm dishes were cultured in Medium D to 60% confluency, then switched to sterol-depleting Medium E for 18-20 hours. Purified, recombinant wild-type, A44P-or D374Y-PCSK9 was added to the medium and cells incubated at 37°C for 4 hours. Cell surface proteins were biotinylated with Sulfo-NHS-SS-Biotin (Campbell Science, Rockford, Illinois, USA), and captured from whole cell extracts using Neutravidin agarose beads (Pierce) as previously described (74). Biotinylated proteins were eluted by boiling in sample buffer supplemented with 5% β-mercaptoethanol, and proteins were resolved on 8% SDS-PAGE and immunoblotted for LDLR and transferrin receptor.

### Circular Dichroism

Custom peptide synthesis (>98% purity) was by BioMatik (Cambridge, Ontario, Canada). Lyophilized peptides were initially dissolved as stock solutions in 0.1% ammonium hydroxide and peptide concentrations were determined by amino acid analysis (SPARC BioCentre, U. of Toronto, Canada) for use in mean residue molar ellipticity calculations. CD spectra were collected on 30 μM samples of peptides in 10 mM Tris, pH 8.5, and 130 mM NaCl with or without 2.5% (w/v) dodecylphosphocholine (DPC) micelles using a Jasco J-815 CD spectropolarimeter at 37°C with a 0.1-cm quartz cuvette. Spectra were acquired from 200 to 250 nm using five accumulations at 20 nm/min and a data integration time of 8 s. Secondary structure deconvolution of spectra was carried out in CDPro with the CONTIN algorithm and SP43 reference set. To measure the apparent affinity of peptide-lipid interactions, peptide (30 μM) was mixed with increasing concentrations of DPC micelles, and CD spectra were collected after each increment as described above.

### Modeling of the PCSK9 N-terminal prodomain

The crystal structure of PCSK9 at 1.9-Å resolution (PDB ID 2QTW)(29) was used to model the missing residues in the N-terminal region of the prodomain. The structure adopted by residues 31 to 60 was built employing the fragment-based de novo protein structure prediction method implemented in ROSETTA v3.8 (40). In preparation for the modeling, 3-mer and 9-mer fragment library files were created with the ROBETTA server (39), identifying protein-fragment structures in the Protein Data Bank that are compatible with the PCSK9 sequence. Based on these libraries, 180,000 refined PCSK9 models including the N-terminal prodomain were built. The known part of the PCSK9 structure was kept fixed during the conformational search. Analysis of the ROSETTA total score against the Cα RMSD relative to the lowest score model was used to evaluate the modeling convergence. The best full-length PCSK9 model was selected as that with the lowest ROSETTA score. Secondary structure prediction with PSIPRED v3.3 (75), as well as protein compactness and local secondary structural features estimated with the “protein meta-structure” approach (76) are consistent with the predicted structure. The final model was analyzed using PyMOL v1.8.4 (Schrödinger, LLC).

### Statistics

Data are presented as the mean ± SEM. Two-sided statistical analysis was determined by One sample t-test or Student’s t-test, as indicated in Figure Legends, using GraphPad Prism 5 software. Data used measurements from at least two distinct sample preparations for each component.

## ACKNOWLEDGMENTS

This work was supported in part by Heart and Stroke Foundation of Ontario Grant G-15-0009352, Canadian Institutes of Health Research (CIHR) Grant 391063 and the Pfizer ASPIRE Program (T.A.L.). A.V-J. thanks FONDECYT research initiation grant #11170223. The Millennium Nucleus of Ion Channels-Associated Diseases (MiNICAD) is a Millennium Nucleus supported by the Iniciativa Científica Milenio of the Ministry of Economy, Development and Tourism (Chile).

## FOOTNOTES

The abbreviations used are: PCSK9, proprotein convertase subtilisin/kexin type 9; LDLR, low density lipoprotein receptor; LDL-C, LDL-cholesterol; Apo, apolipoprotein; FH, familial hypercholesterolemia; GOF, gain-of-function; LOF, loss-of-function; IDR, intrinsically disordered region; AH, amphipathic helix; EGF, epidermal growth factor-like; CHR domain, cysteine-histidine rich domain; CM1, CHR domain module 1

