## Supplemental Data for "A transient amphipathic helix in PCSK9’s prodomain facilitates low-density lipoprotein binding"

Supplemental Figure S1

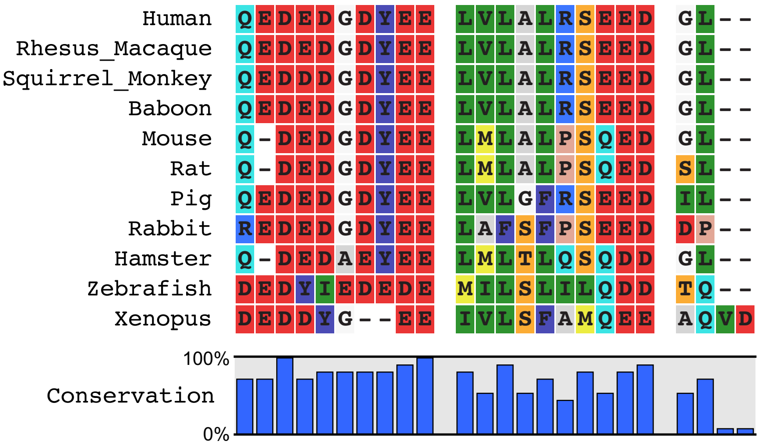


**Supplemental Figure S1.** A multiple sequence alignment of the human PCSK9 aa 31-52 and other vertebrate species highlights evolutionarily conserved regions containing a preponderance of acidic (shaded red) and hydrophobic (shaded green) residues. The RasMol amino color scheme colors amino acids according to traditional amino acid properties (http://life.nthu.edu.tw/~fmhsu/rasframe/COLORS.HTM). Alignment performed using CLC Sequence Viewer v8.0 software (Qiagen Bioinformatics).

Supplemental Figure S2

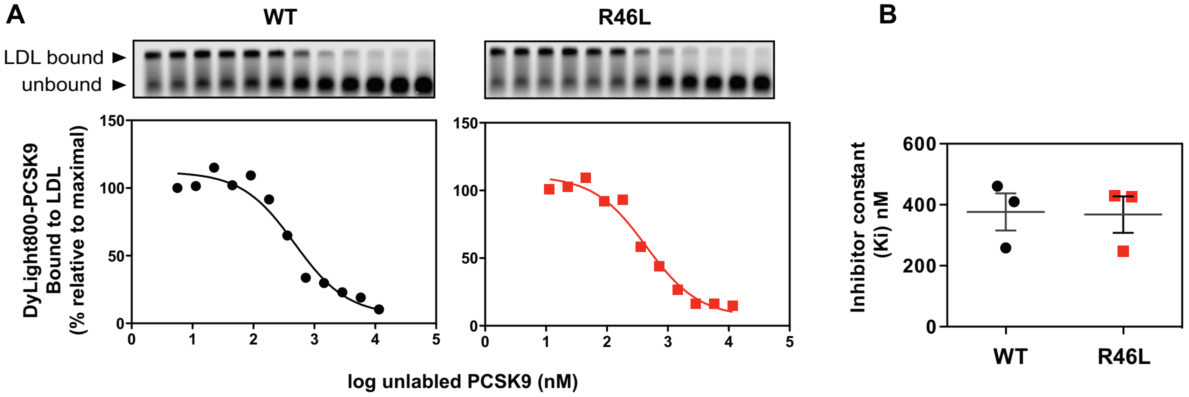


**Supplemental Figure S2.** R46L mutation in PCSK9 does not affect LDL binding affinity. (**A**) In vitro competition binding of WT and R46L forms of PCSK9 to LDL. LDL particles were incubated with Dylight800-labeled PCSK9 in the presence of increasing concentrations of unlabeled competitor proteins. Reaction mixtures were separated on agarose gels (*top*) and fluorophore-labeled PCSK9 binding to LDL was quantified and fitted to competition binding curves using non-linear regression (*bottom*). (**B**) Inhibitor constants (Ki) obtained from curves in *A*. Error bars represent SEM (n=3).

Supplemental Figure S3

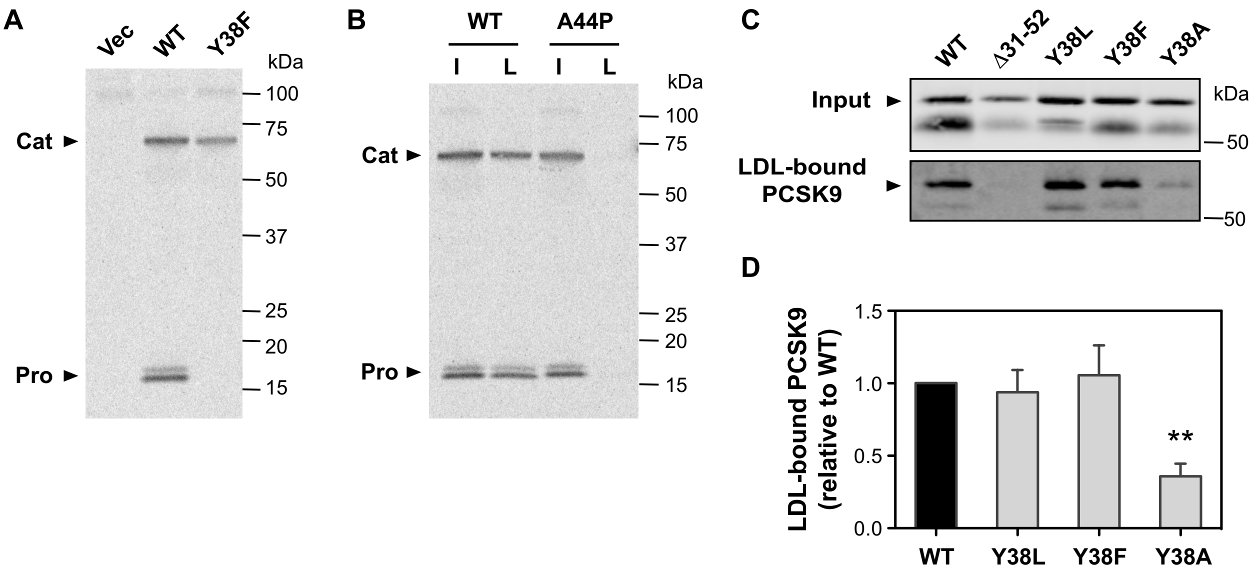


**Supplemental Figure S3.** Role of hydrophobicity at amino acid position 38 (Tyr) in PCSK9-LDL binding. (A) SDS-PAGE analysis of immunoprecipitated wild-type PCSK9 (WT) PCSK9 or Y38F mutant from conditioned medium of transiently transfected HEK293 radiolabeled with $[^{35}S]O_4$. In addition to sulfation of Tyr-38 in the ~16 kDa prodomain, the ~60 kDa catalytic domain/CHR domain segment is also labeled with $[^{35}S]O_4$ on a glycosyl moiety attached to Asn-544 {Benjannet, 2006 #9579}. (B) Analysis of *in vitro* PCSK9-LDL binding reactions. Conditioned medium containing $[^{35}S]$-labeled wild-type PCSK9 (WT) or A44P mutant PCSK9 were incubated with LDL prior to density gradient-ultracentrifugation to isolate an LDL fraction. Lanes are input (I) and LDL-bound PCSK9 (L) (C) Conditioned cell culture medium containing wild-type PCSK9 (WT) or Y38 mutants (Y38L, Y38F or Y38A) were incubated with LDL prior to density gradient-ultracentrifugation and Western blot analysis of LDL-containing fractions. LDL-binding defective Δ31-52 mutant PCSK9 was included as a negative control. (D) Densitometric analyses of western blot in *C*. Error bars represent SEM (n=3). Significant change in LDL binding compared to WT-PCSK9 control (set to 1.0) was determined by One sample t-test: **, $p < 0.01$.

| Mutant | | Primer Sequence (5’→ 3’) |
|---|---|---|
| **Δ33-40** | F: | GCGGGCGCCCGTGCGCAGGAG\|CTGGTGCTAGCCTTGCGT |
| | R: | ACGCAAGGCTAGCACCAG\|CTCCTGCGCACGGGCGCCCGC |
| **Gly-Ser 41-45** | F: | GGACGAGGACGGCGACTACGAGGAG**GGT**G**G**G**AGT**G**G**C**GG**G**A**GTT CCGAGGAGGACGGCCTGGCC |
| | R: | CTCCTCGTAGTCGCCGTCCTCGTCCTCCTGCGCACGGGCGCCCGCG |
| **A44P** | F: | GGAGCTGGTGCTA**C**CCTTGCGTTCCGAGG |
| | R: | CCTCGGAACGCAAGG**G**TAGCACCAGCTCC |
| **L41P** | F: | ACTACGAGGAGC**C**GGTGCTAGCCTTG |
| | R: | CAAGGCTAGCACC**G**GCTCCTCGTAGT |
| **R46L** | F: | GGTGCTAGCCTTGC**T**TTCCGAGGAGGACGG |
| | R: | CCGTCCTCCTCGGAA**A**GCAAGGCTAGCACC |
| **R496W** | F: | GAGTGGGAAGCGG**T**GGGGCGAGCGCATG |
| | R: | CATGCGCTCGCCCC**A**CCGCTTCCCACTC |
| **R469W** | F: | CTCGGGGCCTACA**T**GGATGGCCACAGCCATC |
| | R: | GATGGCTGTGGCCATCC**A**TGTAGGCCCCGAG |
| **F515L** | F: | CCACAACGCTTT**A**GGGGGTGAGGGTG |
| | R: | CACCCTCACCCCC**T**AAAGCGTTGTGG |

**Supplemental Table 1:** Sequences of forward (F) and reverse (R) primers used to mutate C-terminally FLAG-tagged wild-type human PCSK9. Bold, underlined letters indicate mutated base pairs, **|** indicates deletion site.
